# Do Geometric Outliers Identify Important Genes in Single-Cell Foundation Models?

**DOI:** 10.64898/2026.06.22.733850

**Authors:** Justin P. Whalley

## Abstract

Single-cell foundation models (scFMs) produce gene embeddings that are increasingly interpreted biologically, but it is unclear whether different models organise gene space consistently, or whether geometric extremes identify important genes. The same four-metric screen was applied to Geneformer, scGPT and scFoundation, which differ in architecture, training objective and expression encoding. The models agreed weakly on individual outlier genes but showed structured class-level convergence: ribosomal enrichment occurred in all three, although weaker and caller-dependent in scFoundation, while strong mitochondrial enrichment was confined to Geneformer and scGPT. Gene-level agreement was itself uneven, with Geneformer and scGPT overlapping above chance and scGPT and scFoundation not, and caller stability differed between models. Most outliers were not extreme in ESM-2 protein-sequence space, indicating principally model-specific rather than protein-sequence geometry. These patterns did not translate into the tested notions of gene importance: deleting the highest-anomaly Geneformer genes did not impair cell-type annotation beyond matched controls, and covariate-adjusted outlier status was not associated with ClinVar membership in any model. Gene-embedding geometry therefore characterises model-specific structure, but does not provide standalone evidence of downstream leverage or disease relevance.

## 1. Introduction

Single-cell foundation models (scFMs) are increasingly being used to represent transcriptomic data and to support biological inference.^1–4^ These models are commonly assessed through downstream tasks, including cell-type annotation, perturbation prediction and data integration. However, the models also provide gene-level representations that are used to study gene relationships and to nominate genes for experimental follow-up.^1,5,6^ This use assumes that properties of a learned representation carry biological information, rather than reflecting how a particular model was constructed and trained.

The gene representations used by scFMs are produced in different ways. Geneformer uses a BERT-style encoder over rank-ordered gene-token sequences, whereas scGPT combines gene-token and expression-value embeddings within a generative transformer.^1,2,7^ scFoundation uses the xTrimoGene asymmetric encoder–decoder, in which unmasked non-zero positions pass through a larger encoder and continuous expression values are projected through learned auto-discretisation rather than fixed expression bins.^4^ As such, the models differ in their architecture, expression encoding, vocabulary, pretraining objective and training corpus. These differences may affect which genes occupy unusual positions in the learned embedding space.

Most comparative evaluations of scFMs concentrate on downstream performance. Several recent studies have found that foundation models do not consistently outperform simpler approaches in zero-shot evaluation, cell-type annotation or perturbation prediction.^8–10^ Comparatively less attention has been given to whether the gene-embedding geometries produced by different models identify the same genes, or whether geometrically unusual genes have consistent biological properties. This distinction is important because a gene that appears exceptional in one model may not be exceptional in another.

Embedding geometry provides a direct way to make this comparison. Work in natural language processing has described anisotropy, variation in embedding norm and neighbourhood hubness, and has linked some geometrically unusual tokens to unreliable model behaviour.^11–16^ Related geometric analyses have now been proposed for transcriptomic models.^17^ However, geometric extremeness does not by itself demonstrate biological importance or functional consequence. Genes selected using an expression-correlated score may also differ in expression breadth, gene length and functional class, each of which can produce apparently meaningful associations.

The screen used here is deliberately weight-only: it operates on a model’s static gene-embedding matrix without a forward pass or a shared expression dataset. This permits the same geometric calculation to be applied despite incompatible model input encodings and, once protein representations have been computed, to ESM-2 as a non-transcriptomic control. Outcome independence makes the screen useful for model characterisation, not evidence of biological or functional relevance; those claims require the matched and covariate-adjusted tests that follow. The corresponding limitation is that the screen describes input embeddings, not the contextual representations produced for particular cells.

Here, the same four-metric geometric screen is applied to Geneformer, scGPT and scFoundation. These models provide three influential examples of distinct representational designs, but they are not controlled replicates of architecture classes. The comparison is therefore used to establish model-specific differences, rather than to attribute those differences to architecture alone. The resulting outlier sets are tested for robustness to non-parametric re-calling, for recurrence in ESM-2 protein-sequence space,^18^ and for association with ClinVar after adjustment for expression, breadth, length, constraint and gene class. The 50 highest-anomaly Geneformer genes, whose ranking is stable across callers, are then carried into matched deletion experiments in PBMC3k and Tabula Sapiens.

The three models produce substantially different geometric outlier profiles. The Gene-former and scFoundation rankings are largely stable under alternative callers, whereas individual scGPT outlier membership is more sensitive to the calling method. Most outliers do not recur in ESM-2 sequence space, suggesting that their geometry is principally model-specific rather than an intrinsic property of the encoded proteins. In Geneformer, deleting the highest-anomaly genes does not cause greater loss of cell-type annotation performance than deleting matched control genes in either immune dataset. No model’s outlier set shows a detectable association with ClinVar membership after covariate adjustment, and for scGPT a strong unadjusted association does not survive it. Together, these results suggest that embedding geometry is useful for comparing how scFMs represent their vocabularies, but is not stand-alone evidence of functional or disease relevance.

## 2. Methods

### 2.1. Models, embeddings and the geometry screen

Geneformer V2-104M,^1,7^ scGPT^2^ and scFoundation^4^ are audited. The screen is weight-only: it reads a static input gene-embedding matrix and its gene identifiers, and requires no forward pass. It is therefore inexpensive and can be applied wherever a released checkpoint exposes such a matrix. Each gene is scored on four metrics: embedding norm, Euclidean distance to the vocabulary centroid, cosine similarity to the centroid, and mean distance to its *k* = 10 nearest neighbours (isolation). Each is converted to a *z*-score across the vocabulary; a gene is called an outlier if |*z*| *>* 3 on any metric. The metrics are the transcriptomic analogues of quantities studied in NLP embedding spaces: norm and its relation to token statistics,^13^ anisotropy and centroid structure,^11,12^ and neighbourhood irregularity.^14^ Special tokens (`<pad>`, `<mask>`, `<cls>`, `<eos>`) are excluded.

### 2.2. Caller robustness

Outlier identity is re-derived using three robust alternatives to the original |*z*| *>* 3 call, each replacing the mean and standard deviation with the median and the median absolute deviation (MAD): MAD-based *z >* 3, MAD-based *z >* 3.5, and a rank-matched MAD composite. The MAD is scaled by 1.4826, so its thresholds are on the same scale as the original call. Three further callers are retained as diagnostics but excluded from stability assessment: a two-component Gaussian mixture assigns 24.2–49.2% of genes to the minority component across model–metric combinations; percentile 1/99 flags 5.0–6.8% after union across four metrics; and Tukey’s IQR rule (*k* = 3) returns 43 Geneformer outliers and none for the other models. The caller tables and diagnostics supporting each exclusion are provided in the code release.

### 2.3. Non-transcriptomic embedding control

The identical screen is applied to ESM-2 (`esm2_t33_650M_UR50D`) protein embeddings to test whether outliers recur in sequence space.^18^ Genes are mapped to MANE Select v1.3 proteins,^19^ with reviewed UniProt entries as fallback.^20^ Representations are final-layer residue means excluding BOS and EOS; proteins exceeding the 1,022-residue context are embedded in 50%-overlapping windows, averaged first within each window and then, unweighted, across windows. Comparisons use each model’s protein-mappable vocabulary (19,017 genes for Geneformer, 18,748 for scGPT and 18,788 for scFoundation). This control tests recurrence attributable to primary sequence, not functional centrality or biology absent from protein sequence.

### 2.4. Matched-control deletion test

#### Treatment and control sets

The treatment is Geneformer’s 50 highest-anomaly genes, whose ranking was stable across callers (Section 3.2). Two hundred control sets are matched on 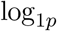 mean expression, expression breadth and 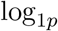 gene length after candidate-pool *z*- scaling,^21^ and exactly on mutually exclusive gene class (Section 2.5). Each control gene is drawn uniformly from the five nearest eligible genes of the same class in that *z*-scaled space. Draws are independent and deduplicated within, but not across, sets. Within each dataset, mean expression was the mean raw count across cells and breadth was the fraction of cells with non-zero expression. The contrast therefore estimates excess deletion cost relative to comparable genes, not arbitrary or low-expression genes.

#### Sensitivity treatment

The procedure is repeated on the 36 genes remaining after removing ribosomal and mitochondrial genes, using a separate 200-draw null matched to that subset.

#### Deletion mechanics

Genes are removed from each cell’s rank-value encoding and sequences are re-padded. This removes a measurement from the model input, approximating an incomplete assay rather than a biological knockout; it does not estimate viability or gene function. Attention masks are derived from token content so ablated sequences cannot attend to padding.

#### Outcome

Geneformer CLS embeddings^7^ are evaluated by macro-*F*_1_ from a multinomial logistic probe with balanced class weights, retrained under five-fold stratified cross-validation.^22^ The treatment is displayed against the central 95% band of the matched-control distribution; inference is one-sided on the lower tail, with empirical *p* = (*b* + 1)*/*(*m* + 1) where *b* of *m* controls are at least as damaging.^23^ Fixed-probe, ARI and NMI results are diagnostic only; the former is in-sample, and the clustering nulls are too wide to resolve an effect of this size, the cluster-ARI band spanning about 43% of baseline against 0.7% for macro-*F*_1_.

#### Datasets

The primary test uses PBMC3k (2,638 cells; 8 cell types).^24,25^ Replication uses a stratified Tabula Sapiens Immune subset:^26,27^ 19,984 cells were subsampled to 4,000 and types with fewer than 10 cells removed, leaving 3,919 cells across 22 of 43 types and 100 matched draws. Each dataset was run under its own pinned environment; both environment fingerprints are recorded.

### 2.5. Annotation panels and class schemes

Ribosomal genes are defined by a pinned 171-gene panel from HGNC groups 728, 729 and 646, matched by Ensembl ID; this includes `RPLP0`–`RPLP2` and `RPSA` while excluding similarly named kinases. Other panels are mitochondrial (`MT-`), loss-of-function constrained, and disease-associated (ClinVar^28^). Constraint uses gnomAD v4.1 pLI *>* 0.9 or LOEUF *<* 0.35.^29^ Using the current LOEUF *<* 0.45 recommendation^30^ gives the same set because all genes between the thresholds already meet the pLI criterion. Panels overlap: 1,784 of 8,285 ClinVar genes (22%) are also constrained.

Every association is therefore evaluated under two schemes. The mutually exclusive scheme assigns one class per gene in the precedence order mitochondrial → ribosomal → constrained → disease → other. The overlapping scheme tests direct membership, allowing a gene to belong to several panels. Reporting both is not redundancy: the two can disagree in direction (Section 3.5).

### 2.6. Statistics, environment and reproducibility

Unadjusted odds ratios use two-sided Fisher tests. Caller-robustness enrichments are Bonferroni-adjusted over four panels; other class-association *p* values are raw and descriptive. ClinVar membership is regressed on outlier status, constraint, 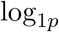 expression, breadth, 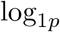 gene length, and ribosomal and mitochondrial indicators. Regression covariates used maximum Human Protein Atlas consensus nTPM across 51 tissues,^31^ the fraction of tissues with nTPM ≥ 1, and the longest Ensembl BioMart gene span. ClinVar membership required at least one pathogenic or likely pathogenic allele in the gene-specific summary dated 4 April 2026. Perfect separation by the mitochondrial indicator requires primary Firth fits with profile-likelihood intervals.^32^ The same universe, covariates and estimator are used for all three models. Matching balance uses standardised mean differences.^33^ Null bands are control 2.5th–97.5th percentiles; empirical *p* values are one-sided. Treatment and controls share a pinned environment; Δ*F*_1_ uses a same-device baseline, whereas 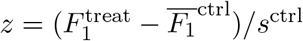 is baseline-invariant.

## 3. Results

### 3.1. Geometric outlier profiles differ substantially among models

Applying the same geometric screen to all three models produced substantially different outlier profiles (Figures 1 and 2). Geneformer yielded 410 outliers across a 20,271-gene vocabulary, scGPT yielded 188 across 60,694 genes, and scFoundation yielded 164 across 19,264 genes. The difference was not limited to the number of outliers. Ribosomal enrichment occurred in all three models, although it was weaker and caller-dependent in scFoundation; strong mitochondrial enrichment was confined to Geneformer and scGPT.

**Fig. 1.**
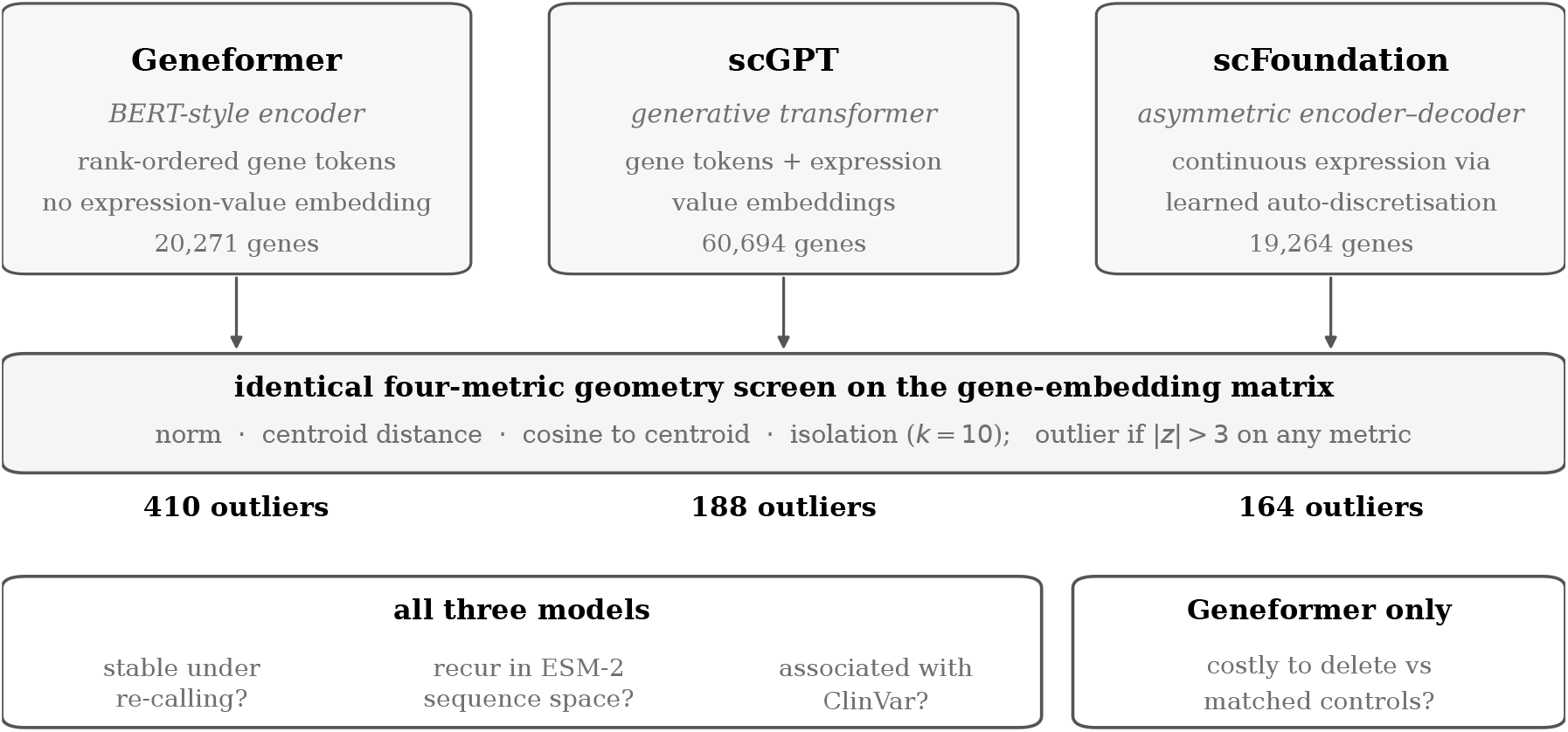
The three models and the shared screen. The models differ in architecture, expression encoding and vocabulary size, and also in pretraining objective, corpus and scale, which are not shown. Caller robustness, ESM-2 recurrence and covariate-adjusted ClinVar association are evaluated for all three; matched deletion is the Geneformer follow-up. With one model per design, the comparison establishes model-specific geometry without attributing it to architecture.

**Fig. 2.**
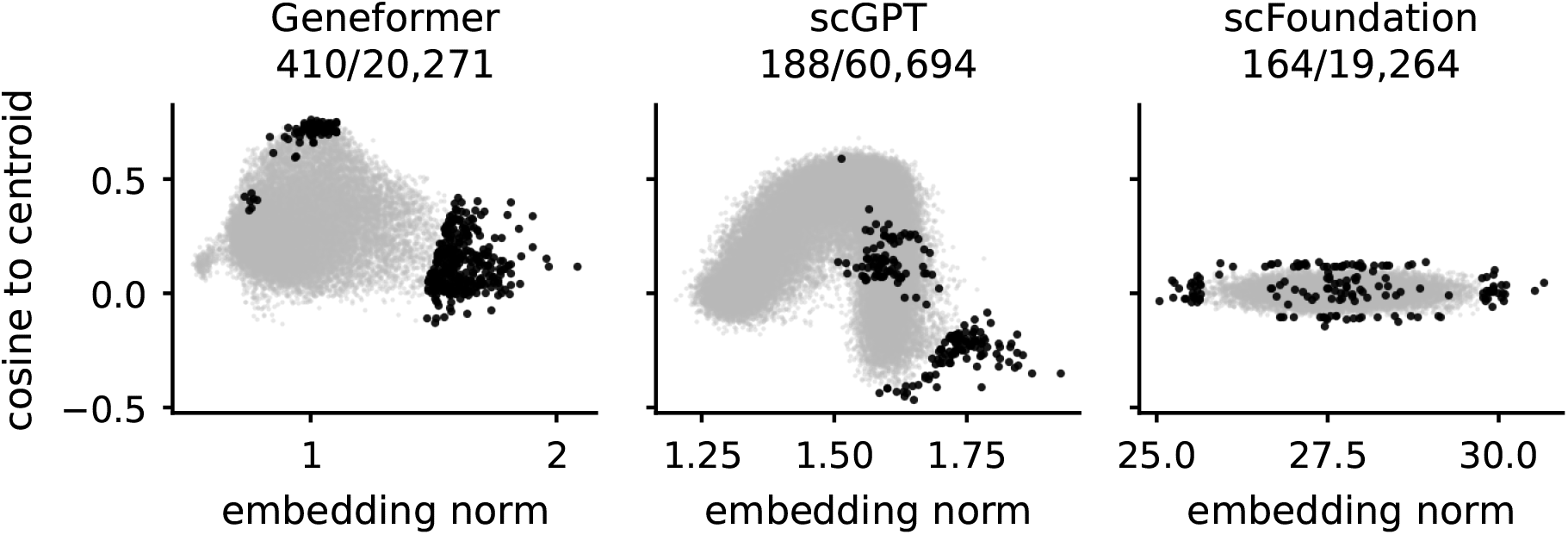
Gene-embedding geometry in each model, with called outliers in black; the cosine axis is shared, the norm axis is model-specific. Geneformer and scGPT each separate into two clouds along the norm axis, the structure behind scGPT’s caller sensitivity (Section 3.2); scFoundation instead forms a narrow band, with cosine to the centroid 0.009*±*0.035 and a 2.5% coefficient of variation in embedding norm.

Counts alone do not establish disagreement, so the outlier sets were compared directly on the 18,915 genes shared by all three vocabularies (Table 1). Agreement is graded rather than uniform. Geneformer and scGPT share 72 outliers against 2.1 expected by chance, a 34-fold excess (hypergeometric *p* = 1 *×* 10^−99^), yet their Jaccard index is only 0.172 and the genomewide correlation between their anomaly scores is *ρ* = 0.186. Geneformer and scFoundation share 8 against 3.3 expected (*p* = 0.018, and 0.055 after Bonferroni correction across the three pairwise tests, so nominal only), and scGPT and scFoundation share a single gene against 0.9 expected (*p* = 0.58), which is chance. One gene is an outlier in all three. The two models that represent genes as discrete tokens agree more with each other than either does with the continuous-expression model, while still agreeing only weakly in absolute terms. That ordering is consistent with expression encoding mattering, but it is not evidence for it: corpus, objective, vocabulary and scale differ alongside, and one model per design cannot separate them.

**Table 1.** Direct cross-model agreement on the 18,915 genes shared by all three vocabularies. Hypergeometric overlap *p* values are raw across three pairwise tests.

| Model pair | Overlap | Expected | Jaccard | $\rho$ | Raw overlap $p$ |
| --- | --- | --- | --- | --- | --- |
| Geneformer, scGPT | 72 | 2.1 | 0.172 | 0.186 | $1 \times 10^{-99}$ |
| Geneformer, scFoundation | 8 | 3.3 | 0.015 | 0.026 | 0.018 |
| scGPT, scFoundation | 1 | 0.9 | 0.004 | 0.007 | 0.58 |

Over the same 18,915 shared genes, class composition also differed. Mitochondrial enrichment was 163-fold in Geneformer and 455-fold in scGPT, whereas scFoundation had no mitochondrial outliers at all; ribosomal enrichment was 44.8-, 18.9and 4.4-fold across the same three models. Expression separated the models in the same way: median maximum HPA expression among outliers was 19 times the non-outlier median in Geneformer and 12 times in scGPT, but slightly below it in scFoundation. Geneformer outliers were also enriched for loss-of-function-constrained genes (OR 1.46 under the mutually exclusive scheme), and its 50 highest-anomaly genes had much higher median maximum HPA expression than the vocabulary overall (1,813 against 42 nTPM), despite heterogeneous expression breadth. Several of the highest-ranked genes were tissue-restricted. Downstream effects could therefore reflect expression and class rather than geometry.

### 3.2. Caller stability differs among models

Because the original calls use |*z*| *>* 3 on metrics whose distributions are skewed or bimodal, outlier sets were re-derived under robust alternatives (Section 2.2). For Geneformer, MAD-based recalling recovers every one of the 410 original |*z*| *>* 3 outliers, so containment, the fraction of that original set recovered, is 100%; the rank ordering is preserved (*ρ* = 0.956) and the top-50 genes are identical. scFoundation behaves similarly (containment 97%, *ρ* = 0.999, top-50 overlap 49/50).

Geneformer’s Jaccard index against the MAD set is nonetheless only 0.492, a consequence of asymmetric caller breadth rather than loss of the original core set: MAD calling recovers all 410 original outliers and adds a further 423, so the original set is a strict subset and the union in the denominator is inflated. The original calls are a conservative core.

Containment must be read against its ceiling, since a stricter caller returning fewer genes cannot contain them all; Figure 3 marks these ceilings. This matters most for scFoundation under MAD *z >* 3.5, where containment falls to 16%. That caller returns only 27 genes and all 27 are among the original 164, so the bar sits exactly on its ceiling. As such, the low value reflects the breadth of the caller rather than instability of the outlier set. scGPT is the exception: containment is 83% and top-50 overlap 22/50, so individual gene identity is method-sensitive, a consequence of its bimodal norm distribution, under which mean- and median-based standardisation weight the tails differently. Class-level associations are nevertheless stable across the MAD family for Geneformer and scGPT alike; no class significant under the original call loses significance.

**Fig. 3.**
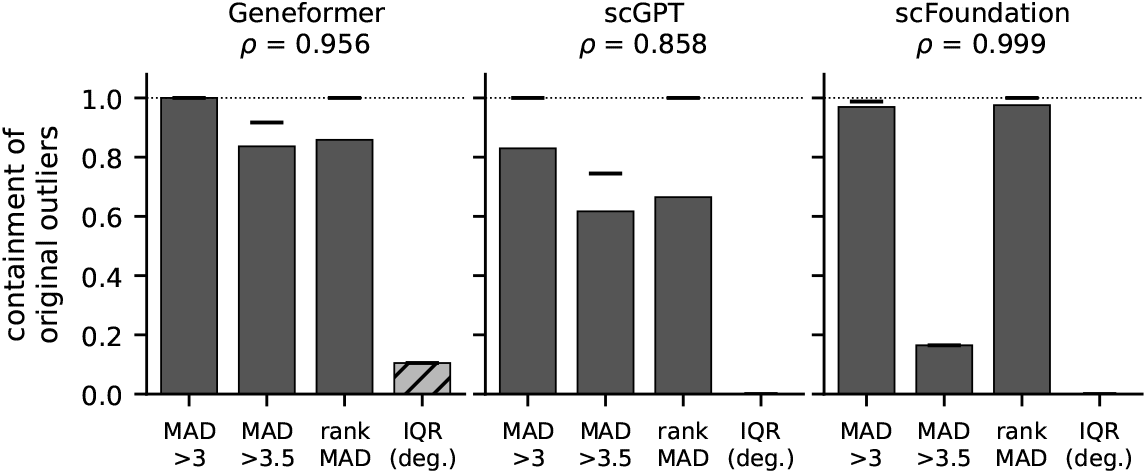
Outlier identity under alternative callers. Bars give containment, the fraction of original |*z*| *>* 3 outliers recovered; ticks mark the maximum attainable containment min(*n*_new_, *n*_orig_)*/n*_orig_, so a bar meeting its tick is saturated, not unstable. The hatched IQR bar is one of three degenerate callers reported but excluded (Section 2.2). *ρ* is the Spearman correlation between original and MAD-based anomaly scores.

### 3.3. Outliers are largely model-specific, with a smaller sequence-related component

If geometric outliers reflected properties of the encoded proteins rather than of a particular model, the same screen applied to protein-sequence embeddings should recover them (Section 2.3). Across 19,017 shared genes, Geneformer and ESM-2 outlier sets had Jaccard 0.054 and anomaly-score correlation *ρ* = 0.072 (averaged over all three scFMs, 0.023 and 0.042). Forty-nine genes overlapped against 11.7 expected (*p <* 10^−15^), but only four appeared in both top-50 lists. Thus most geometry was model-specific, with a smaller sequence-related component.

The shared component included eight mitochondrially encoded, 22 ribosomal and seven olfactory-receptor genes. Under mutually exclusive classes (Figure 4), all 82 constrained and 79 of 82 disease-associated outliers were scFM-specific, as were 52 of 74 ribosomal outliers. Mitochondrial genes were the exception, with only 2 of 10 scFM-specific because the other 8 recurred in ESM-2. Their distinct genetic code^34^ and hydrophobic membrane proteins^35^ make this consistent with sensitivity to genuine sequence distinctiveness rather than failure to recover any class.

**Fig. 4.**
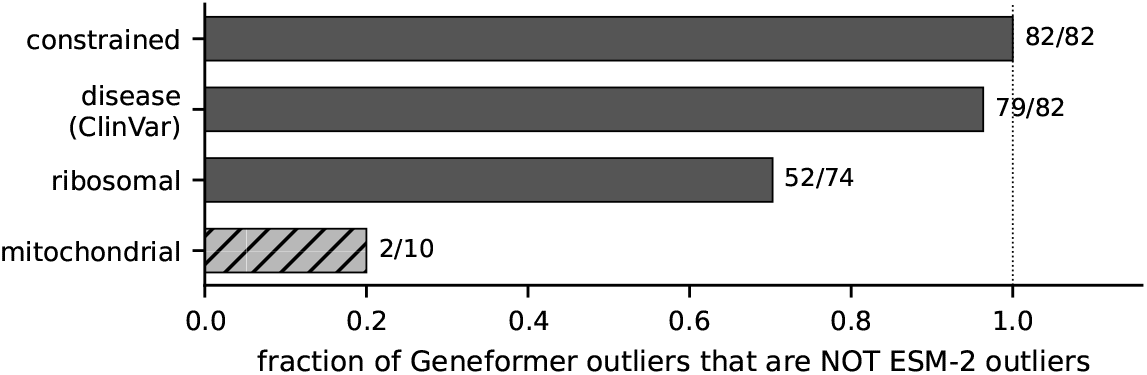
Fraction of Geneformer outliers that do not recur in ESM-2, by mutually exclusive class over 19,017 shared genes; bars are labelled with the scFM-specific count. Constrained, disease-associated and ribosomal outliers are largely scFM-specific. The hatched mitochondrial bar is short because only 2 of its 10 outliers are scFM-specific, the other 8 being ESM-2 outliers as well.

### 3.4. Deleting outliers did not exceed the matched-control null

If geometric extremeness marked genes on which the model depends, deleting them should cost more cell-type annotation accuracy than deleting comparable genes. From a PBMC3k baseline of 0.924, deleting the 50 highest-anomaly genes left macro-*F*_1_ 0.00280 below the mean of 200 matched controls (*z* = −1.50). Twelve controls were at least as damaging (*p* = 0.065), placing the treatment inside the empirical 95% null band (Figure 5).

**Fig. 5.**
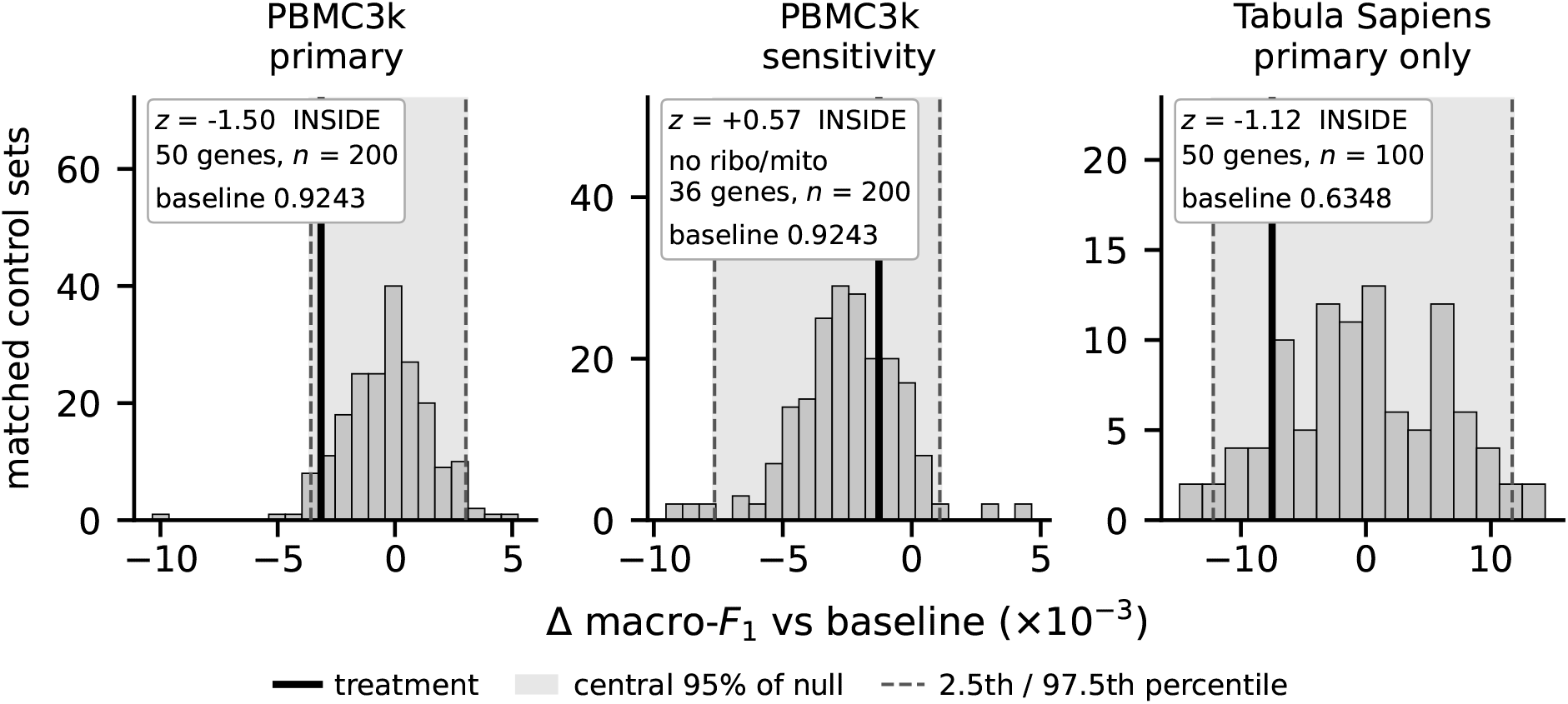
Matched-control deletion results. Each panel gives Δ macro-*F*_1_ for gene sets matched to the treatment on expression level, breadth, gene length and class; shading is the 2.5th–97.5th percentile null band and the vertical line is the treatment. Axes are not shared and each panel states its own baseline and draw count, so the three histograms are not a common null. Left, the 50 highest-anomaly genes in PBMC3k (*z* = −1.50). Centre, the 36 remaining after removing ribosomal and mitochondrial genes, against their own null (*z* = +0.57). Right, the primary arm on Tabula Sapiens (*z* = −1.12); its sensitivity arm was not replicated. All three fall inside their bands. Cluster ARI is excluded (Section 2.4).

Ribosomal and mitochondrial genes account for 28% of the treatment set, so any cost might follow from removing those classes rather than from geometry. The 36 genes remaining once they were excluded sat 0.00122 above their own control mean (*z* = +0.57; 145 of 200 controls at least as damaging, *p* = 0.73). The weak degradation seen in the full treatment was not observed in the non-ribosomal, non-mitochondrial sensitivity arm; because each arm was assessed against its own matched null, the analysis does not test the difference between arms.

#### Replication on a less saturated task

PBMC3k annotation is close to saturated at 0.924, leaving little room for a deletion to register. The same treatment was therefore applied to a Tabula Sapiens immune subset with a baseline of 0.635, where a real cost would have more room to appear. It left macro-*F*_1_ 0.0073 below the mean of 100 controls, again inside the null (*z* = −1.12; 14 controls at least as damaging, *p* = 0.149). Token exposure was comparable between treatment and controls, so the result is not explained by the treatment genes appearing less often in the analysed cells; matching was exact on class though imperfect on expression (SMD 0.283) and length (0.112). The sensitivity arm was run on PBMC3k only. Agreement across two datasets with very different baselines supports the primary result, though it does not show the effect to be exactly zero.

### 3.5. No model’s outliers show a detectable adjusted ClinVar association

Geometric outliers are enriched for functionally central gene classes, which invites the inference that they are disease-relevant. ClinVar^28^ membership was therefore regressed on outlier status, adjusting for expression, breadth, length, constraint and gene class (Section 2.6). Across 18,911 complete cases, none of the three outlier sets showed a detectable association (Table 2a). Geneformer gave an adjusted OR of 1.05 (95% confidence interval, CI, 0.82–1.34, *p* = 0.72), and scFoundation 0.98 (0.69–1.38, *p* = 0.91). The scGPT result shows what the adjustment does to a strong unadjusted signal: its unadjusted OR of 2.95 (*p* = 1.8 *×* 10^−7^) fell to 1.21 (0.76–1.98, *p* = 0.42) under the same adjustment. The intervals remain compatible with effects in either direction, including a nearly twofold association for scGPT, so these are nulls of detection rather than demonstrations of exact independence.

**Table 2.** ClinVar association over 18,911 complete cases. GF is Geneformer. **(a)** covariate-adjusted. **(b)** unadjusted, under two annotation schemes, which estimate different quantities. Non-mitochondrial sensitivity fits and the full specification grid are in the code release.

| **(a) Adjusted** |  |  |  |  |
| --- | --- | --- | --- | --- |
| Tier | Gene set | Unadj. OR | Adjusted OR (95% CI) | *p* |
| PRIMARY (Firth) | Geneformer ($n = 388$) | 1.16 | 1.05 (0.82–1.34) | 0.72 |
| PRIMARY (Firth) | scGPT ($n = 102$) | 2.95 | 1.21 (0.76–1.98) | 0.42 |
| PRIMARY (Firth) | scFoundation ($n = 161$) | 0.80 | 0.98 (0.69–1.38) | 0.91 |
| PRIMARY (Firth) | GF ∩ scGPT (exploratory; $n = 72$) | 3.59 | 1.44 (0.79–2.69) | 0.23 |

| **(b) Unadjusted, by annotation scheme** |  |  |  |
| --- | --- | --- | --- |
| Gene set | Class scheme | OR | *p* |
| GF ∩ scGPT ($n = 72$) | Overlapping | 3.59 | $5.0 \times 10^{-7}$ |
| GF ∩ scGPT ($n = 72$) | Mutually exclusive | 0.385 | $1.6 \times 10^{-3}$ |
| All Geneformer ($n = 388$) | Overlapping | 1.16 | 0.15 |
| All Geneformer ($n = 388$) | Mutually exclusive | 0.519 | $3.8 \times 10^{-8}$ |

The 72-gene Geneformer–scGPT subset was similarly enriched before adjustment (OR 3.59, *p* = 5.0*×*10^−7^), but not afterwards (1.44, 0.79–2.69, *p* = 0.23). It is reported as exploratory because it was selected post hoc, and its attenuation cannot be assigned to a single covariate without nested models.

That same unadjusted enrichment is also sensitive to something other than covariates. Under direct membership the 72 shared outliers give a ClinVar odds ratio of 3.59 (*p* = 5.0 *×* 10^−7^); under the mutually exclusive scheme the same 72 genes give 0.385 (*p* = 1.6 *×* 10^−3^), an apparent depletion (Table 2b). The reversal follows from the precedence order. Constrained genes are labelled before disease genes, 22% of ClinVar genes are also constrained, and these outliers are enriched for constraint at OR 7.36, so the residual disease category is stripped of the genes that would have carried the enrichment. Both odds ratios are correctly computed and answer different questions. A gene-class enrichment reported without its scheme can therefore point either way.

## 4. Discussion

The same screen returned sets that differed in size, class composition and caller stability, so a finding in one scFM should not be assumed to transfer to another. The discrete-token models agreed most, but architecture, encoding, vocabulary, objective, corpus and scale all varied, with one model per design. This ordering motivates controlled comparison; it does not identify its cause or establish that one design is preferable.

In Geneformer and scGPT, outliers had much higher maximum tissue expression and were enriched for functionally central classes; scFoundation showed much less of both, with outlier expression slightly below the vocabulary median. Expression breadth did not follow the same pattern, running lower among Geneformer outliers and higher among scGPT’s. Pretraining token frequencies were not measured, so it cannot be shown that these are the genes each model encountered most often, and expression measured in a held-out dataset is only a proxy for exposure during training. Gene expression is heavy-tailed,^36^ and the observed enrichment toward the head of that distribution is an inference rather than a demonstration. This is still useful information about a model: it describes how learned gene representations are distributed across a vocabulary, and it differs markedly between the models examined here.

Deleting the highest-anomaly Geneformer genes did not degrade cell-type annotation more than matched controls in either dataset. Separately, none of the three models’ outlier sets showed a detectable adjusted ClinVar association; scGPT provides the clearest example, with its odds ratio falling from 2.95 to 1.21. Genes selected on an expression-correlated criterion can therefore appear functionally central while showing no detectable additional signal for the outcomes tested here, cell-type annotation under matched deletion and covariate-adjusted ClinVar membership. That is the distinction matched or covariate-adjusted controls are there to make.

The absence of a detectable adjusted association is itself an interesting result. The outlier sets are strongly enriched for genes a biologist would call central, ribosomal, mitochondrial and constrained, yet none of the three models showed a residual link to genes already curated as disease-associated once expression, breadth, length, constraint and class were accounted for. Their centrality is of a kind already visible without the model. An expression-correlated score will preferentially select highly expressed genes and, through correlated covariates, genes over-represented in constraint and disease panels, so an unmatched enrichment cannot show that the score itself carries information. The annotation-scheme result makes the same point from another direction: overlapping and mutually exclusive panels estimate different quantities and can point in opposite directions, so a gene-class enrichment should state its scheme and report both where they disagree.

The transferable contribution is therefore the audit design rather than a universal list of outlier genes: a checkpoint-level score generates a hypothesis, and matched or covariate-adjusted analyses test whether it adds information for a specified outcome. The outlier sets reported here are specific to three checkpoints and should not be assumed to transfer. The screen half of that design is also cheap in a way that matters as these models proliferate. When a released checkpoint exposes a static gene-embedding matrix and identifiers, the screen needs no evaluation data and can be run immediately, raising no question of evaluation-set contamination. Downstream benchmarking cannot avoid that question, since it requires data whose overlap with the pretraining corpus is rarely documented. Inexpensive checks on what a model has learned are worth having as validated computational models take a larger role alongside experimental new-approach methodologies.^37^

### Limitations

This work concentrates on three well-established models chosen for their different representational designs, and their geometries differ enough that conclusions about another model would need the same analysis repeated rather than assumed.^38^ Space restricted the biological tests to immune datasets and cell-type annotation, though nothing in the design prevents the same test on other tissues, or on perturbation prediction,^39^ network inference or atlas integration.^40^ Within those tests, Tabula Sapiens uses a filtered 22-class subset with 100 draws and no sensitivity arm, and matching remains imperfect (expression SMD 0.32 in PBMC3k and 0.28 in Tabula Sapiens; length 0.11 in Tabula Sapiens).^33^ The screen also describes geometry without defining what a well-behaved embedding space would look like, and reads static input embeddings rather than contextual representations computed for individual cells. Finally, the pretraining corpora are too large to work with here, so why particular genes occupy extreme positions cannot be established. Work on language models has traced anomalous tokens to under-exposure during training,^15,16^ and the equivalent account for scFMs would need that access.

## 5. Conclusion

Geneformer, scGPT and scFoundation produce substantially different gene-embedding geometries, and genes exceptional in one model are largely not exceptional in another. Most are also not outliers in protein-sequence space, suggesting that the geometry principally reflects how each model represents its vocabulary rather than an intrinsic gene property. The genes these screens return are central ones, but on the two outcomes tested that centrality was not information the model had added: it was already visible from expression and gene class, and it did not extend to excess loss of cell-type annotation under matched deletion or to ClinVar association after adjustment. For a model that exposes a static gene-embedding matrix and its gene identifiers, the screen is cheap, requiring no forward pass or expression data, and can be applied as soon as the checkpoint is released. That, rather than any particular list of genes, is what carries forward.

## Supporting information

Supplemental Table 1: per-gene geometry and annotation table

## Data and code availability

Code, versioned inputs, saved outputs and a traceability table linking the reported analyses to their generating scripts and outputs are available at: https://github.com/jpwhalley/matched-null-audit The submission release is tagged `psb-2027-submission` at commit `90798e87d133`. The software environment and input checksums required for reproduction are provided there. Supplemental Table 1, posted with this preprint, is the per-gene annotation table underlying Tables 1 and 2, identical to `data/Table S1.csv` in that release.

## References

1. C. V. Theodoris, L. Xiao, A. Chopra, M. D. Chaffin, Z. R. Al Sayed, M. C. Hill, H. Mantineo, E. M. Brydon, Z. Zeng, X. S. Liu and P. T. Ellinor, Transfer learning enables predictions in network biology, Nature 618, 616 (2023).

2. H. Cui, C. Wang, H. Maan, K. Pang, F. Luo, N. Duan and B. Wang, scGPT: toward building a foundation model for single-cell multi-omics using generative AI, Nature Methods 21, 1470 (2024).

3. F. Yang, W. Wang, F. Wang, Y. Fang, D. Tang, J. Huang, H. Lu and J. Yao, scBERT as a large-scale pretrained deep language model for cell type annotation of single-cell RNA-seq data, Nature Machine Intelligence 4, 852 (2022).

4. M. Hao, J. Gong, X. Zeng, C. Liu, Y. Guo, X. Cheng, T. Wang, J. Ma, X. Zhang and L. Song, Large-scale foundation model on single-cell transcriptomics, Nature Methods 21, 1481 (2024).

5. F. Pedrocchi, F. Barkmann, A. Joudaki and V. Boeva, Sparse autoencoders reveal interpretable features in single-cell foundation models, bioRxiv (2025), bioRxiv 2025.10.22.681631; latest revision 16 June 2026.

6. I. Kendiukhov, Sparse autoencoders reveal organized biological knowledge but minimal regulatory logic in single-cell foundation models: a comparative atlas of Geneformer and scGPT, arXiv (2026), arXiv:2603.02952 [q-bio.GN].

7. H. Chen, M. S. Venkatesh, J. Gómez Ortega, S. V. Mahesh, T. N. Nandi, R. K. Madduri, K. Pelka and C. V. Theodoris, Scaling and quantization of large-scale foundation model enables resource-efficient predictions in network biology, Nature Computational Science 6, 450 (2026).

8. R. Boiarsky, N. M. Singh, A. Buendia, A. P. Amini, G. Getz and D. Sontag, Deeper evaluation of a single-cell foundation model, Nature Machine Intelligence 6, 1443 (2024).

9. K. Z. Kedzierska, L. Crawford, A. P. Amini and A. X. Lu, Zero-shot evaluation reveals limitations of single-cell foundation models, Genome Biology 26, p. 101 (2025).

10. C. Ahlmann-Eltze, W. Huber and S. Anders, Deep-learning-based gene perturbation effect prediction does not yet outperform simple linear baselines, Nature Methods 22, 1657 (2025).

11. K. Ethayarajh, How contextual are contextualized word representations? Comparing the geometry of BERT, ELMo, and GPT-2 embeddings, in Proceedings of the 2019 Conference on Empirical Methods in Natural Language Processing and the 9th International Joint Conference on Natural Language Processing (EMNLP-IJCNLP), eds. K. Inui, J. Jiang, V. Ng and X. Wan (Association for Computational Linguistics, Hong Kong, China, November 2019).

12. J. Mu and P. Viswanath, All-but-the-top: Simple and effective postprocessing for word representations, in International Conference on Learning Representations (ICLR), (ICLR, Vancouver, Canada, 2018).

13. M. Oyama, S. Yokoi and H. Shimodaira, Norm of word embedding encodes information gain, in Proceedings of the 2023 Conference on Empirical Methods in Natural Language Processing, eds. H. Bouamor, J. Pino and K. Bali (Association for Computational Linguistics, Singapore, December 2023).

14. M. Radovanović, A. Nanopoulos and M. Ivanović, Hubs in space: Popular nearest neighbors in high-dimensional data, Journal of Machine Learning Research 11, 2487 (2010).

15. J. Rumbelow and M. Watkins, SolidGoldMagikarp (plus, prompt generation) (2023), Blog post, LessWrong / AI Alignment Forum.

16. S. Land and M. Bartolo, Fishing for Magikarp: automatically detecting under-trained tokens in large language models, in Proceedings of the 2024 Conference on Empirical Methods in Natural Language Processing, eds. Y. Al-Onaizan, M. Bansal and Y.-N. Chen (Association for Computational Linguistics, Miami, Florida, USA, November 2024).

17. W. Gilpin, The cell as a token: high-dimensional geometry in language models and cell embeddings, Bioinformatics 41, p. btaf595 (2025).

18. Z. Lin, H. Akin, R. Rao, B. Hie, Z. Zhu, W. Lu, N. Smetanin, R. Verkuil, O. Kabeli, Y. Shmueli, dos Santos Costa, M. Fazel-Zarandi, T. Sercu, S. Candido and A. Rives, Evolutionary-scale prediction of atomic-level protein structure with a language model, Science 379, 1123 (2023).

19. J. Morales, S. Pujar, J. E. Loveland, A. Astashyn, R. Bennett, A. Berry, E. Cox, C. Davidson, O. Ermolaeva, C. M. Farrell, R. Fatima, L. Gil, T. Goldfarb, J. M. Gonzalez, D. Haddad, M. Hardy, T. Hunt, J. Jackson, V. S. Joardar, M. Kay, V. K. Kodali, K. M. McGarvey, A. McMahon, J. M. Mudge, D. N. Murphy, M. R. Murphy, B. Rajput, S. H. Rangwala, L. D. Riddick, F. Thibaud-Nissen, G. Threadgold, A. R. Vatsan, C. Wallin, D. Webb, P. Flicek, E. Birney, K. D. Pruitt, A. Frankish, F. Cunningham and T. D. Murphy, A joint NCBI and EMBL-EBI transcript set for clinical genomics and research, Nature 604, 310 (2022).

20. The UniProt Consortium, UniProt: the Universal Protein Knowledgebase in 2023, Nucleic Acids Research 51, D523 (2023).

21. E. A. Stuart, Matching methods for causal inference: a review and a look forward, Statistical Science 25, 1 (2010).

22. F. Pedregosa, G. Varoquaux, A. Gramfort, V. Michel, B. Thirion, O. Grisel, M. Blondel, P. Prettenhofer, R. Weiss, V. Dubourg, J. Vanderplas, A. Passos, D. Cournapeau, M. Brucher, M. Perrot and É. Duchesnay, Scikit-learn: machine learning in Python, Journal of Machine Learning Research 12, 2825 (2011).

23. B. Phipson and G. K. Smyth, Permutation P-values should never be zero: calculating exact Pvalues when permutations are randomly drawn, Statistical Applications in Genetics and Molecular Biology 9, p. Article 39 (2010).

24. G. X. Y. Zheng, J. M. Terry, P. Belgrader, P. Ryvkin, Z. W. Bent, R. Wilson, S. B. Ziraldo, T. D. Wheeler, G. P. McDermott, J. Zhu, M. T. Gregory, J. Shuga, L. Montesclaros, J. G. Underwood, D. A. Masquelier, S. Y. Nishimura, M. Schnall-Levin, P. W. Wyatt, C. M. Hindson, R. Bharadwaj, A. Wong, K. D. Ness, L. W. Beppu, H. J. Deeg, C. McFarland, K. R. Loeb, W. J. Valente, N. G. Ericson, E. A. Stevens, J. P. Radich, T. S. Mikkelsen, B. J. Hindson and J. H. Bielas, Massively parallel digital transcriptional profiling of single cells, Nature Communications 8, p. 14049 (2017).

25. F. A. Wolf, P. Angerer and F. J. Theis, SCANPY: large-scale single-cell gene expression data analysis, Genome Biology 19, p. 15 (2018).

26. The Tabula Sapiens Consortium, The Tabula Sapiens: a multiple-organ, single-cell transcriptomic atlas of humans, Science 376, p. eabl4896 (2022).

27. CZI Cell Science Program, S. Abdulla, B. Aevermann, P. Assis, S. Badajoz, S. M. Bell, E. Bezzi, B. Cakir, J. Chaffer, S. Chambers, J. M. Cherry, T. Chi, J. Chien, L. Dorman, P. Garcia-Nieto, N. Gloria, M. Hastie, D. Hegeman, J. Hilton, T. Huang, A. Infeld, A.-M. Istrate, I. Jelic, K. Katsuya, Y. J. Kim, K. Liang, M. Lin, M. Lombardo, B. Marshall, B. Martin, F. McDade, C. Megill, N. Patel, A. Predeus, B. Raymor, B. Robatmili, D. Rogers, E. Rutherford, D. Sadgat, A. Shin, C. Small, T. Smith, P. Sridharan, A. Tarashansky, N. Tavares, H. Thomas, A. Tolopko, M. Urisko, J. Yan, G. Yeretssian, J. Zamanian, A. Mani, J. Cool and A. Carr, CZ CELLxGENE Discover: a single-cell data platform for scalable exploration, analysis and modeling of aggregated data, Nucleic Acids Research 53, D886 (2025).

28. M. J. Landrum, J. M. Lee, M. Benson, G. R. Brown, C. Chao, S. Chitipiralla, B. Gu, J. Hart, D. Hoffman, W. Jang, K. Karapetyan, K. Katz, C. Liu, Z. Maddipatla, A. Malheiro, K. McDaniel, M. Ovetsky, G. Riley, G. Zhou, J. B. Holmes, B. L. Kattman and D. R. Maglott, ClinVar: improving access to variant interpretations and supporting evidence, Nucleic Acids Research 46, D1062 (2018).

29. K. J. Karczewski, L. C. Francioli, G. Tiao, B. B. Cummings, J. Alföldi, Q. Wang, R. L. Collins, K. M. Laricchia, A. Ganna, D. P. Birnbaum, L. D. Gauthier, H. Brand, M. Solomonson, N. A. Watts, D. Rhodes, M. Singer-Berk, E. M. England, E. G. Seaby, J. A. Kosmicki, R. K. Walters, K. Tashman, Y. Farjoun, E. Banks, T. Poterba, A. Wang, C. Seed, N. Whiffin, J. X. Chong, K. E. Samocha, E. Pierce-Hoffman, Z. Zappala, A. H. O’Donnell-Luria, E. V. Minikel, B. Weisburd, M. Lek, J. S. Ware, C. Vittal, I. M. Armean, L. Bergelson, K. Cibulskis, K. M. Connolly, M. Covarrubias, S. Donnelly, S. Ferriera, S. Gabriel, J. Gentry, N. Gupta, T. Jeandet, D. Kaplan, C. Llanwarne, R. Munshi, S. Novod, N. Petrillo, D. Roazen, V. Ruano-Rubio, A. Saltzman, M. Schleicher, J. Soto, K. Tibbetts, C. Tolonen, G. Wade, M. E. Talkowski, C. A. Aguilar Salinas, T. Ahmad, C. M. Albert, D. Ardissino, G. Atzmon, J. Barnard, L. Beaugerie, E. J. Benjamin, M. Boehnke, L. L. Bonnycastle, E. P. Bottinger, D. W. Bowden, M. J. Bown, J. C. Chambers, J. C. Chan, D. Chasman, J. Cho, M. K. Chung, B. Cohen, A. Correa, D. Dabelea, M. J. Daly, D. Darbar, R. Duggirala, J. Dupuis, P. T. Ellinor, R. Elosua, J. Erdmann, T. Esko, M. Färkkilä, J. Florez, A. Franke, G. Getz, B. Glaser, S. J. Glatt, D. Goldstein, C. Gonzalez, L. Groop, C. Haiman, C. Hanis, M. Harms, M. Hiltunen, M. M. Holi, C. M. Hultman, M. Kallela, J. Kaprio, S. Kathiresan, B.-J. Kim, Y. J. Kim, G. Kirov, J. Kooner, S. Koskinen, H. M. Krumholz, S. Kugathasan, S. H. Kwak, M. Laakso, T. Lehtimäki, R. J. F. Loos, S. A. Lubitz, R. C. W. Ma, D. G. MacArthur, J. Marrugat, K. M. Mattila, S. McCarroll, M. I. McCarthy, D. McGovern, R. McPherson, J. B. Meigs, O. Melander, A. Metspalu, B. M. Neale, P. M. Nilsson, M. C. O’Donovan, D. Ongur, L. Orozco, M. J. Owen, C. N. A. Palmer, A. Palotie, K. S. Park, C. Pato, A. E. Pulver, N. Rahman, A. M. Remes, J. D. Rioux, S. Ripatti, D. M. Roden, D. Saleheen, V. Salomaa, N. J. Samani, J. Scharf, H. Schunkert, M. B. Shoemaker, P. Sklar, H. Soininen, H. Sokol, T. Spector, P. F. Sullivan, J. Suvisaari, E. S. Tai, Y. Y. Teo, T. Tiinamaija, M. Tsuang, D. Turner, T. Tusie-Luna, E. Vartiainen, M. P. Vawter, J. S. Ware, H. Watkins, R. K. Weersma, M. Wessman, J. G. Wilson, R. J. Xavier, B. M. Neale, M. J. Daly, D. G. MacArthur and G. A. D. Consortium, The mutational constraint spectrum quantified from variation in 141,456 humans, Nature 581, 434 (2020).

30. K. Chao, J. Goodrich, P. Darnowsky, R. Panchal, K. Laricchia, M. Wilson and K. Samocha, gnomAD v4.1.1 (2026), gnomAD browser news. Recommends LOEUF < 0.45 for v4.1.1.

31. M. Uhlén, L. Fagerberg, B. M. Hallström, C. Lindskog, P. Oksvold, A. Mardinoglu, Å. Sivertsson, C. Kampf, E. Sjöstedt, A. Asplund, I. Olsson, K. Edlund, E. Lundberg, S. Navani, C. A.-K. Szigyarto, J. Odeberg, D. Djureinovic, J. O. Takanen, S. Hober, T. Alm, P.-H. Edqvist, H. Berling, H. Tegel, J. Mulder, J. Rockberg, P. Nilsson, J. M. Schwenk, M. Hamsten, K. von Feilitzen, M. Forsberg, L. Persson, F. Johansson, M. Zwahlen, G. von Heijne, J. Nielsen and F. Pontén, Tissue-based map of the human proteome, Science 347, p. 1260419 (2015).

32. D. Firth, Bias reduction of maximum likelihood estimates, Biometrika 80, 27 (1993).

33. P. C. Austin, Balance diagnostics for comparing the distribution of baseline covariates between treatment groups in propensity-score matched samples, Statistics in Medicine 28, 3083 (2009).

34. S. Anderson, A. T. Bankier, B. G. Barrell, M. H. L. de Bruijn, A. R. Coulson, J. Drouin, I. C. Eperon, D. P. Nierlich, B. A. Roe, F. Sanger, P. H. Schreier, A. J. H. Smith, R. Staden and I. G. Young, Sequence and organization of the human mitochondrial genome, Nature 290, 457 (1981).

35. R. Eaglesfield and K. Tokatlidis, Targeting and insertion of membrane proteins in mitochondria, Frontiers in Cell and Developmental Biology 9, p. 803205 (2021).

36. C. Furusawa and K. Kaneko, Zipf’s law in gene expression, Physical Review Letters 90, p. 088102 (2003).

37. U.S. Food and Drug Administration, Roadmap to reducing animal testing in preclinical safety studies, tech. rep., U.S. Food and Drug Administration (Washington, DC, 2025).

38. P. Qiu, Q. Chen, H. Qin, S. Fang, Y. Zhang, Y. Zhang, T. Xia, L. Cao, Y. Zhang, X. Fang, Y. Li and L. Hu, BioLLM: A standardized framework for integrating and benchmarking single-cell foundation models, Patterns 6, p. 101326 (2025).

39. J. M. Replogle, R. A. Saunders, A. N. Pogson, J. A. Hussmann, A. Lenail, A. Guna, L. Mascibroda, E. J. Wagner, K. Adelman, G. Lithwick-Yanai, N. Iremadze, F. Oberstrass, D. Lipson, J. L. Bonnar, M. Jost, T. M. Norman and J. S. Weissman, Mapping information-rich genotypephenotype landscapes with genome-scale Perturb-seq, Cell 185, 2559 (2022).

40. M. D. Luecken, M. Büttner, K. Chaichoompu, A. Danese, M. Interlandi, M. F. Mueller, D. C. Strobl, L. Zappia, M. Dugas, M. Colomé-Tatché and F. J. Theis, Benchmarking atlas-level data integration in single-cell genomics, Nature Methods 19, 41 (2022).

